# Cancer type classification in liquid biopsies based on sparse mutational profiles enabled through data augmentation and integration

**DOI:** 10.1101/2021.03.09.434391

**Authors:** Alexandra Danyi, Myrthe Jager, Jeroen de Ridder

## Abstract

Identifying the cell of origin of cancer is important to guide treatment decisions. However, in patients with ‘cancer of unknown primary’ (CUP), standard diagnostic tools often fail to identify the primary tumor. As an alternative, machine learning approaches have been proposed to classify the cell of origin based on somatic mutation profiles in the genome of solid tissue biopsies. However, solid biopsies can cause complications and certain tumors are not accessible. A promising alternative would be liquid biopsies, which contain ctDNA originating from the tumor. Problematically, somatic mutation profiles of tumors obtained from liquid biopsies are inherently extremely sparse and current machine learning models fail to perform in this setting.

Here we propose an improved machine learning method to deal with the sparse nature of liquid biopsy data. Firstly, we downsample the SNVs in the samples in order to mimic sparse data conditions. Then extensive data augmentation is performed to artificially increase the number of training samples in order to enhance model robustness under sparse data conditions. Finally, we employ data integration to merge information from i) somatic single nucleotide variant (SNV) density across the genome, ii) somatic SNVs in driver genes and iii) trinucleotide motifs. Our adapted method achieves an average accuracy of 0.88 on the data where only 70% of SNVs are retained, which is comparable to an average accuracy of 0.87 with the original model on the full SNV data. Even when only 2% of the data is retained, the average accuracy is 0.65 compared to 0.41 with the original model. The method and results presented here open the way for application of machine learning in the detection of the cell of origin of cancer from sparse liquid biopsy data.

**Author Summary:** The identification of the ‘cell of origin’ of cancer is an important step towards more personalized cancer care, but this remains a challenge for patients with ‘cancer of unknown primary’ (CUP) where the source of the malignancy cannot be identified even after extensive clinical assessment with standard diagnostic methods. Somatic mutation profile-based ‘cell of origin’ classification has emerged in recent years as a promising alternative diagnostic tool that could circumvent the issues of standard CUP diagnostic. In this approach the somatic mutations are obtained from whole genome sequencing (WGS) of solid tissue biopsies from the tumor. However, needle biopsies from tumor tissue can be challenging, as accessibility to the tumor can be limited and taking a biopsy can cause further complications. For these reasons, liquid biopsies have been proposed as a safer alternative to solid tissue biopsies. Problematically, the circulating tumor DNA fragments available in e.g. blood typically represent a much scarcer tumor source than conventional solid tissue biopsies and therefore liquid biopsies give rise to sparse somatic mutation profiles. Therefore it is crucial to investigate the applicability of sparse somatic mutation profiles in the identification of ‘cell of origin’ and explore potential improvements of the data analysis and prediction models to overcome sparsity.

## Introduction

Identification of the ‘cell of origin’ of cancer - which is representative of its anatomical origin and histological characteristics - is an important step in effective cancer treatment, as it has been established that the cell of origin can be a strong predictor of response to therapy and overall prognosis [1, 2]. Importantly, tailoring targeted treatments to the cell of origin can result in more successful treatment and better prognosis for ‘cancer of unknown primary’ (CUP) patients, where the cell of origin is often difficult to determine with standard histopathology techniques [3]. One particularly promising alternative to identify the cell of origin in CUP patients is based on somatic mutations derived from whole genome sequencing (WGS) data. Recent studies demonstrate proof of concept of this, by leveraging machine learning to accurately identify the cell of origin using somatic mutation density profiles obtained in solid biopsies of primary and metastatic cancers [4, 5]. However, in many cases taking solid tissue biopsies is challenging, as accessibility to the tumor can be limited. Moreover, conventional needle biopsies are invasive and may lead to clinical complications in one out of six procedures [6]. Even in case a biopsy is feasible, solid tissue biopsies cannot be used for early detection and screening since the location of the tumor is not yet known by definition.

Blood plasma of cancer patients contains circulating tumor DNA (ctDNA) which can be used to detect somatic mutations [7,8,9]. Liquid biopsies of the blood may, therefore, provide a valuable alternative to solid biopsies. Moreover, conventional solid tissue biopsy can only capture tumor heterogeneity to a limited extent as only one spatial location of the tumor is sampled, while ctDNA originates from the whole tumor and thus captures more of the tumor heterogeneity [10, 11]. A major challenge for ctDNA-based cancer diagnostics, however, is that ctDNA only represents a small fraction of the total cell-free DNA (cfDNA). As a result, the amount of tumor DNA that can be captured in liquid biopsies is much lower than in conventional solid tissue biopsies. While in patients with advanced-stage cancers the ctDNA concentration can exceed 10% of the total cfDNA (thus increasing the chance of detecting comprehensive somatic mutation profiles), it is significantly lower in patients with cancer types such as glioma, medulloblastoma, bladder, renal and gastroesophageal cancer [10]. Moreover, ctDNA levels are likely to be lower in early-stage tumors where the tumor burden is low [12, 7]. While ctDNA levels can vary depending on tumor type and disease stage, the generally limited amount of ctDNA in liquid biopsies results in a much sparser somatic mutation profile when compared to solid tissue biopsies.

Liquid biopsies are only a viable alternative to solid biopsies if there is a certain level of concordance between the somatic mutations obtained from ctDNA and the tissue, which is generally high in case of metastatic tumors. According to a prospective study on advanced metastatic cancers, 72% of somatic SNV/indel mutations found in solid tissue biopsy samples were detected by the applied liquid biopsy assay with high-intensity sequencing [13] and similarly in an earlier study on metastatic prostate and breast cancers, the average concordance was determined as 88% [14]. However, in a recent study on stage I-IV non-small lung cell cancer (NSCLC) patients, the percentage of concordant mutations between cfDNA and tissue biopsy ranged between only 12.4% (stage I) and 73.8% (stage IV) [15]. This shows that lower ctDNA concentrations in early stage cancer can result in a more dramatic sparsity which might complicate the utilization of somatic mutations to identify the cell of origin, while metastatic cancers with higher tumor load are less affected by this issue. Therefore, there is no generally defined concordance level of somatic mutations in a liquid biopsy when compared to a corresponding solid biopsy sample, but it is clear that the variable concordance of somatic mutations poses a challenge to diagnostic applications due to potentially high levels of sparsity.

Despite the sparsity of somatic mutation profiles in liquid biopsies, efforts have already been made to utilize liquid biopsy data for the identification of the cell of origin of cancer. The majority of these studies successfully apply targeted approaches [7, 16], which mainly focus on mutation density in cancer driver genes as input features for cancer type classification. Due to the low abundance of ctDNA, many driver mutations can be missed due to sampling artifacts in WGS, therefore targeted sequencing is necessary to ensure detection sensitivity. However, adding genome-wide information could improve the classification. According to a recent study on genome-wide liquid biopsies of postoperative early stage residual cancers, the integration of genome-wide mutation data allowed sensitive residual disease detection by overcoming the limitations of sparsity [17]. Additionally, previous work on WGS data from solid tissue biopsies have already established that somatic SNV density at 1 Mb scale is the most prominent predictor of cancer type as it represents the genomic imprint of the cell of origin chromatin organization, with passenger somatic SNVs being the most prominent contributors [4, 5]. However, the performance of a support vector machine classifier based on passenger SNVs was considerably reduced on artificially generated sparse data [4]. This indicates that current state-of-the-art classifiers might not be robust enough to reliably predict tumor type from somatic mutation profiles which are extremely sparse, such as those obtained from liquid biopsies.

In our work, we explore the utilization of sparse genome-wide somatic mutation data in the classification of the cell of origin of cancer. High quality genome-wide somatic mutation data obtained from ctDNA is very scarce [17], and by no means sufficient to support the training of robust classifiers. Therefore sparse SNV samples are generated based on WGS of primary cancer samples from the PCAWG dataset [18] to model ctDNA conditions. Then, in order to address the challenge of sparsity and enable model robustness, we propose to improve on a state-of-the-art classification approach in two ways. First of all, we recognize that, in a sparse somatic data setting, massive training set sizes are required to enable the classifier to become robust against feature sparsity (i.e. missing somatic mutations), therefore we apply extensive data augmentation. Secondly, to complement the genome-wide somatic SNV density features, we propose to leverage data integration of two additional features derived from the SNV data: trinucleotide counts and mutation density on driver genes. Mutational signatures, which differ between cancer types and reflect the result of diverse oncogenic processes, are constructed from the whole mutation spectra that consists of trinucleotide (e.g.: ACA>AGA) mutation frequencies [19]. These trinucleotide features were utilized for cancer type classification earlier [5]. Trinucleotide mutational contexts are composite features and therefore can be less affected by scarcely available SNV mutations. Additionally, while frequently mutated driver genes can represent one of the major hallmarks of cancer and are commonly shared between cancer types [20], certain combinations of affected driver genes can differ between different tissues [21], which has been used as a basis for cancer type classification before [7] and driver gene mutation density features were found to have some predictive power in a few cancer types (for instance Panc-AdenoCA) [5].

Our results demonstrate that both data augmentation as well as data integration considerably increase the robustness of the classifier. As a result, when the original solid tissue biopsy samples are downsampled to retain only 70% of somatic mutations (an approximation of general sparsity in liquid biopsy of late stage and metastatic cancers [13, 15]), the deep learning model proposed here reaches classification accuracies similar to previous state-of-the-art classifiers on solid biopsy (i.e. non-sparse) WGS mutation profiles. These findings raise the possibility that somatic mutation profiles derived from liquid biopsies can aid the detection of the cell of origin in CUP patients.

## Results

### Assessment of state-of-the-art deep learning model on sparse somatic mutation data

We first assessed the classification performance of a state-of-the-art deep learning model proposed in Jiao et al. 2020 in a liquid biopsy setting (**Materials and methods**). This model is based on genome-wide SNV density features across 2,897 bins where each bin represents a 1 Mb region of the genome. Sparse mutation profiles were artificially generated based on the PCAWG data collection [18], a dataset that consists of 2,374 primary cancer samples across 16 cancer types with Soft-Messarc having the smallest sample size of 33 samples and Liver-HCC representing the biggest cancer type with 304 samples (see **S1 Table** for tumor type abbreviations). To model the somatic mutation sparsity of liquid biopsy samples, sparse samples were obtained by selecting a subset of all somatic SNVs (mutation downsampling) from each PCAWG sample. Multiple sparse datasets were generated by retaining 70%, 25%, 10%, 5% and 2% of all somatic SNVs (**Fig 1a**). The first measurement was designated at 70%, as this value is an approximation of the reported percentage of somatic mutations that can be detected in liquid biopsies compared to solid tissue biopsies in most advanced and metastatic cancers [13, 15]. Lower downsampling levels were chosen to mimic lower ctDNA concentrations, which have been observed in certain cancer types and in case of low tumor burden [7, 10]. The full somatic SNV data of all 2,374 samples was also classified in order to have baseline accuracies of the original classification setting from [5] for each cancer type.

**Fig 1.**
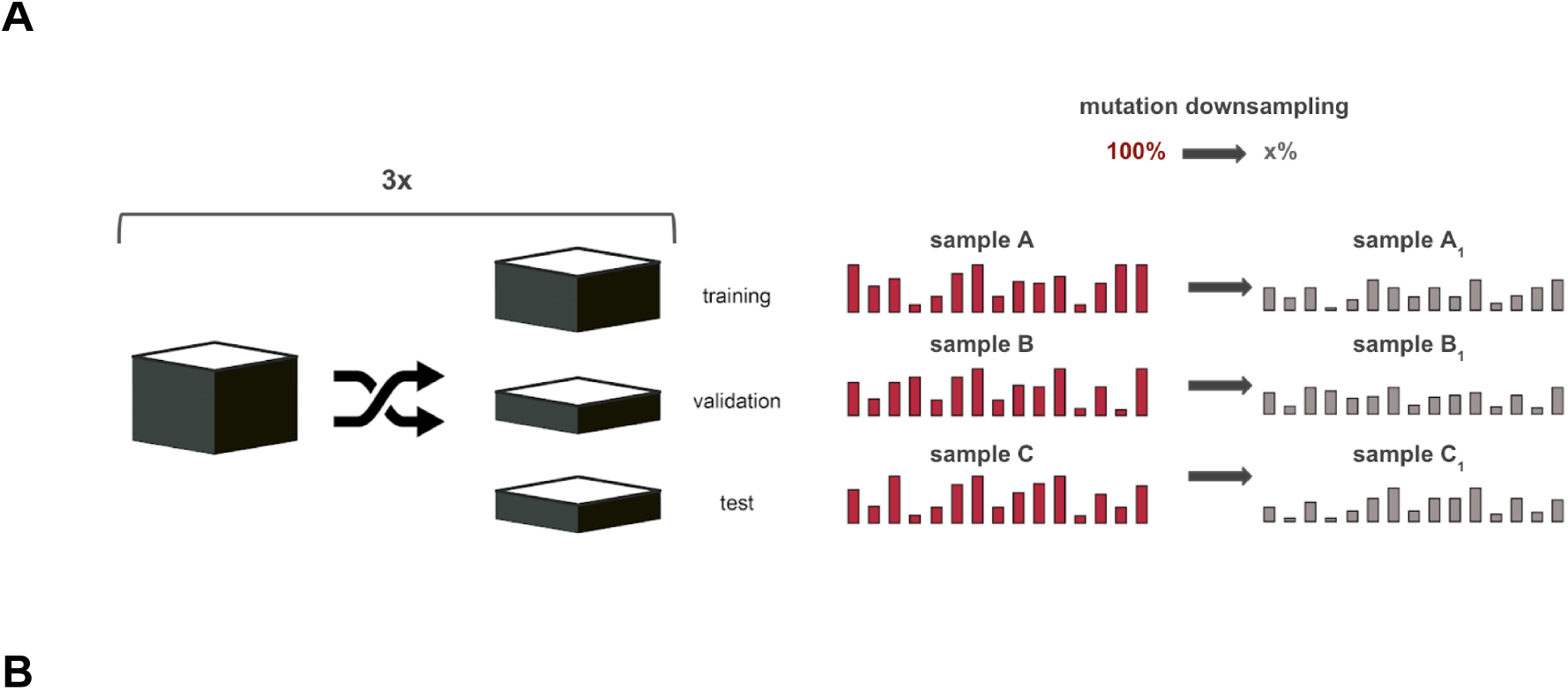
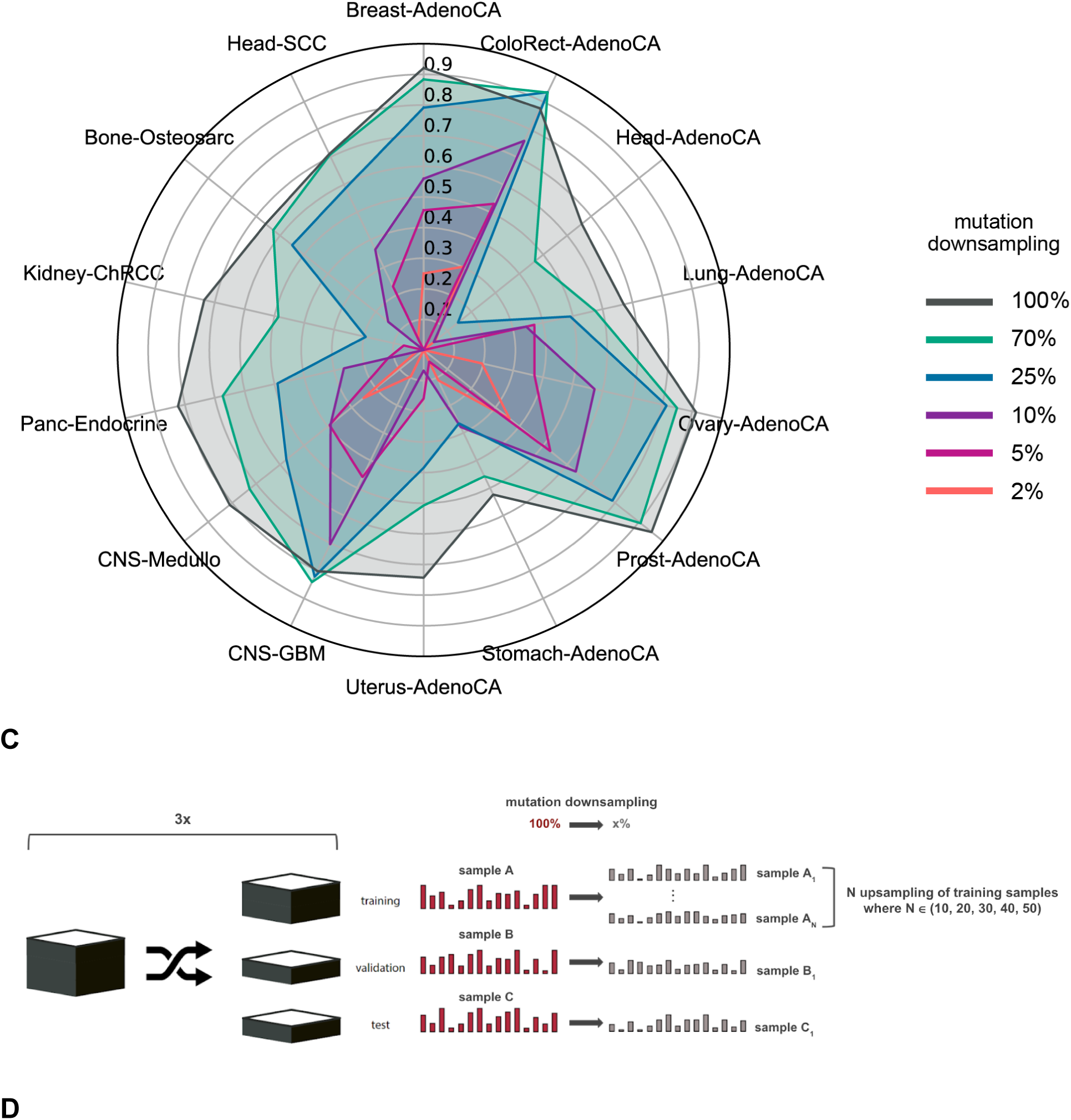
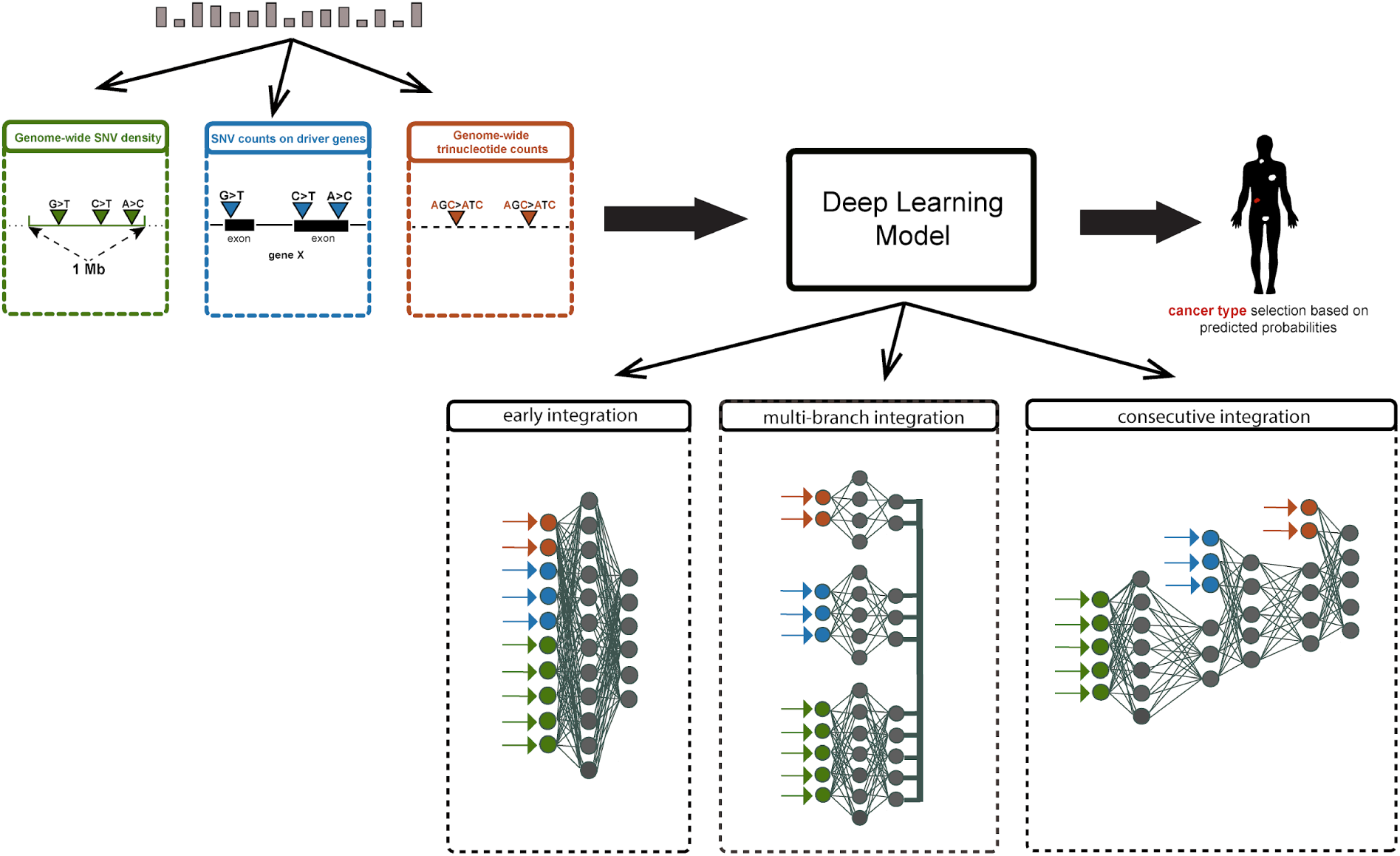
State-of-the-art model performance on the generated sparse datasets and overview of proposed model improvements. **(A)** Schematic overview of sparse somatic SNV data generation. Sparse samples were generated by retaining 70%, 25%, 10%, 5% and 2% of somatic mutations from each original sample (mutation downsampling) for the training, validation and test sets. Results were evaluated in 3-fold stratified shuffle-split cross-validation. **(B)** Cancer types classification accuracies of somatic SNV data with different (70%, 25%, 10%, 5%, 2%) mutation downsampling rates and full somatic SNV data (100%). The axes represent the accuracy for the cancer types, expressed in average F1 scores (stratified shuffle split cross-validation with three partitions). Only cancer types with the lowest sparse data accuracies are shown. **(C)** Schematic overview of data augmentation. Sparse somatic SNV samples (subsamples of each original WGS sample) were generated by retaining 70%, 25%, 10%, 5% and 2% of SNVs from each original sample (mutation downsampling). Data augmentation was applied solely on the training set, where instead of 1 subsample, N=(10, 20, 30, 40, 50) different subsamples were generated based on each full somatic SNV sample (sample upsampling). Results were evaluated in 3-fold shuffle-split cross-validation. **(D)** Schematic overview of data integration of trinucleotide motifs and driver gene mutation densities to tackle sparse somatic SNV data. The integration of the three feature sets was tested in different deep learning architectures: early integration, multi-branch integration and consecutive integration.

As an initial test, predictions were made on the sparse datasets with the original model trained on the full (100%) somatic SNV data. As expected, accuracy dropped dramatically in case of each sparse dataset, except at 70% (**S1 Fig**). When the model is trained and tested on the full 100% SNV data, our results are in line with previous findings [5] and show that for all but five cancer types the classification accuracy exceeds 0.7, and for most cancer types is well above 0.8 (**S1-2 Fig, Fig 1b**). Stomach-AdenoCA is the worst performing tumor type and is often misclassified as Panc-AdenoCA or Eso-AdenoCA due to similar biological origin, as described in the earlier study [5].

In the next step, we aimed to assess the effect of SNV downsampling on the training procedure and retrained the original model on each sparse dataset, instead of applying the model trained on the full data throughout all sparse test datasets. Overall, when the model is trained and tested on sparse data, a better accuracy trend is appreciable compared to the results when the model is trained on the full somatic SNV data but tested on sparse data (**S1-2 Fig, Fig 1b**). Generally, when the downsampling rate increases, classification accuracy decreases, often substantially. When only 2% of somatic SNVs are retained, an average accuracy below 0.2 in 10 out of 16 tumor types was observed (**S2 Fig**). Still, for six of the cancer types a classification accuracy exceeding 0.5 was retained, suggesting that even at this sparsity level some signal is retained that can be used by the classifier. Interestingly, for ColoRect-AdenoCA the classification accuracy improved in the 70% downsampled mutation experiment compared to the 100% baseline (**Fig 1b**). This might suggest that downsampling the mutations has a regularization effect on the model for this particular cancer type. Similarly, the average prediction accuracy of CNS-nonDiffGBM and Lung-SCC also slightly improved at 70% (**S2 Fig**).

In summary, the classification accuracy gradually drops as the number of retained somatic mutations decreases, with the least but still considerable performance decrease at 70%, even if the classification model was trained on the corresponding sparse data set (**Fig 1b, S2 Fig**). This indicates that the current model is not suitable for liquid biopsy data that has a sparser somatic profile than conventional solid tissue biopsies.

### Data augmentation

Data augmentation is a technique that was introduced in the field of deep learning to aid image classification problems where the available training data is limited. Essentially, data augmentation enhances the available data in terms of size, for instance through basic sample (image) manipulations (geometrical or color transformations, random erasing, image mixing etc.) or model-based augmentation methods (GANs, adversarial training etc.) [22]. Data augmentation addresses the issue of overfitting during the training process, as the augmented data represents a much more comprehensive collection of possible data points. Therefore, better model generalization and robustness can be achieved. By applying different data augmentation approaches, the error rate of the model can be reduced considerably, even up to 12.1% [23,24,25,26].

Contrary to traditional machine learning techniques, the performance of deep learning models keeps improving as the training data size is increasing, according to a power-law relationship [27]. Hence, we hypothesized that reduced classification performance due to sparsity of somatic mutation profiles obtained from liquid biopsies can be mitigated by substantially increasing the training set size. By providing the classifier with more and diverse training data it should have more information to learn robust feature patterns under sparse data conditions. However, since genome-wide somatic mutation data from liquid biopsy of more than a few samples per cancer type is not yet readily available, we investigated if the concept of ‘in silico’ data augmentation can be applied. To achieve sufficiently large training set sizes, augmented training samples were generated based on multiple random downsampling of somatic SNVs in PCAWG WGS samples (**Fig 1c**).

Multiple data augmentation experiments (generating N subsamples from each original sample, where N∈(10,20,30,40,50); see section ‘Data augmentation’ in **Materials and methods**) were performed at each mutation downsampling rate in order to identify a potential inflexion point after which further data augmentation does not improve the performance (**Fig 1c**). The different data augmentation rates were compared based on average accuracy (average ratio of correctly classified samples in stratified shuffle split cross-validation with three partitions). For all downsampling rates, 10x data augmentation greatly improves the performance of the model to an average accuracy of 0.51 compared to 2% baseline with 0.41, 0.66 compared to the 5% baseline with 0.56, 0.73 compared to the 10% baseline with 0.63, 0.81 compared to 25% baseline with 0.75 and 0.87 compared to the 70% baseline with 0.83, respectively (**Fig 2a**). At the lowest mutation downsampling rates (when 25% and 70% of all SNVs were retained), performance tops with using only 20x data augmentation, at an average accuracy of 0.84 compared to 25% baseline with 0.75 and 0.88 compared to the 70% baseline with 0.83, respectively. At the highest mutation downsampling rates (when retaining only 2% and 5% of all SNVs), the application of 50x data augmentation shows the most considerable average performance improvement compared to the baseline experiments using the original model without data augmentation, at an average accuracy of 0.57 compared to 2% baseline with 0.41 and 0.69 compared to the 5% baseline with 0.56, respectively (**Fig 2a**).

**Fig 2.**
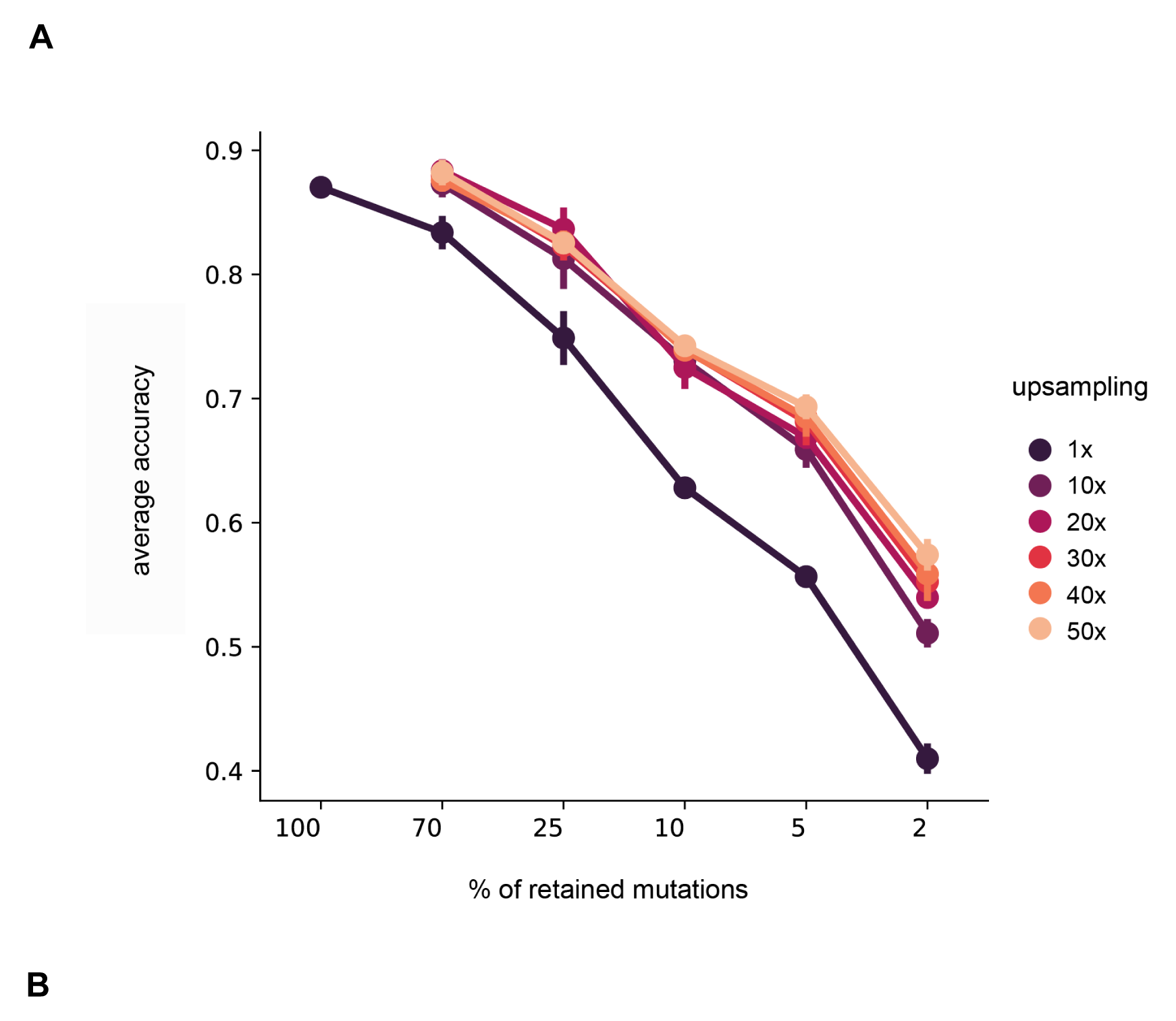
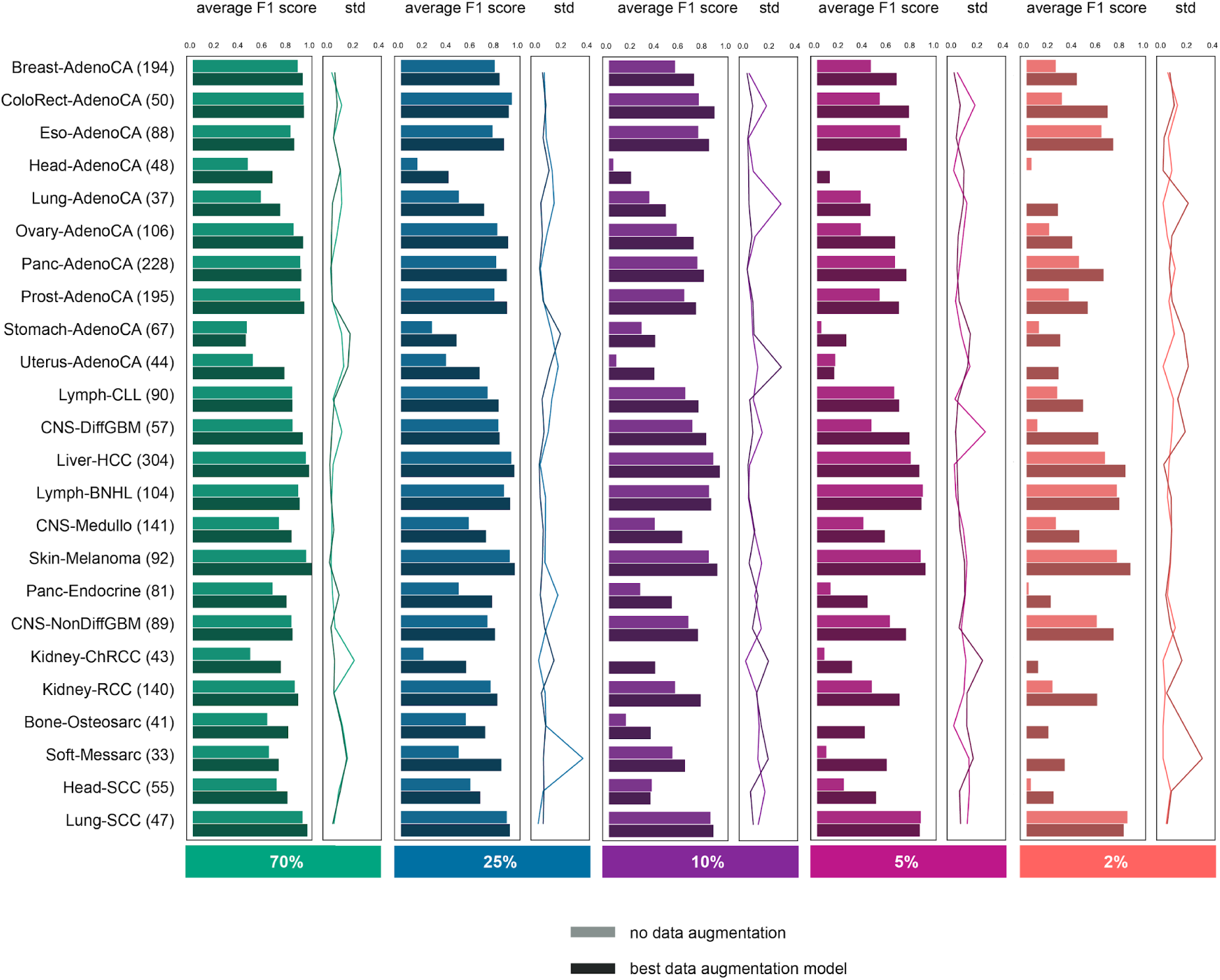
Classifier performance with data augmentation. **(A)** Overall effect of data augmentation at different mutation downsampling rates (70%, 25%, 10%, 5%, 2%). Average accuracy (average % of accurately classified samples in 3 folds) is shown at different downsampling rates with and without applied data augmentation setups **(B)** Classifier performance at different mutation downsampling rates with the best data augmentation setup, expressed in F1-scores per cancer type (average and std is shown based on 3 folds). The best data augmentation setup was selected based on average accuracy values: 70% - 20x, 25% - 20x, 10% - 30x, 5% - 50x, 2% - 50x.

The model performance was also assessed at cancer type level in all mutation downsampling experiments (expressed in average F1 scores) (**Fig 2b**, **S3 Fig**). When only 70% and 25% of somatic mutations are retained, data augmentation improves the classifier performance for all cancer types, except ColoRect-AdenoCA, while at 10% mutation downsampling only the accuracy of Head-SCC does not improve. At higher downsampling rates (retaining only 2% and 5% of SNVs) data augmentation improves the classifier performance for all cancer types, except for Uterus-AdenoCA, Lymph-BNHL and Lung-SCC at 5% and Head-AdenoCA and Lung-SCC at 2%. While Lymph-BNHL and Lung-SCC are both classified with high accuracy across all downsampling rates even without data augmentation, Uterus-AdenoCA and Head-AdenoCA show a rather dramatic drop in classification accuracy as the level of sparsity increases and contrary to other cancer types, data augmentation remained ineffective at 2% and 5% downsampling for these two types (**S2 Fig, Fig 2b**). Lower classification accuracy can also be partially associated with lower sample numbers in a few cancer types (**Fig 2b**). When retaining only 2% and 5% of somatic SNVs, all samples are persistently misclassified from a few cancer types without data augmentation (**Fig 2b**). These results clearly show that data augmentation is fundamental when dealing with classification of the cell of origin of cancer from sparse genome-wide mutation data and it can considerably improve the classification accuracy even at higher downsampling rates (2%, 5%, 10%).

### Data integration

To compensate for the higher data sparsity, integrating additional genomic features based on the SNV data (i.e. genome-wide SNV density bins, mutation density on driver genes and genome-wide trinucleotide motifs) may improve classification accuracy even further. Model design is a crucial element in this process, even more so when multiple feature types have to be processed in the model. We therefore assessed three main architecture types for the integration of the three SNV-derived features: early integration, multi-branch integration and consecutive integration. The aim of testing these main architecture types was to assess different feature integration approaches, where consecutive integration represents a hierarchical feature integration approach, multi-branch integration applies feature integration in the hidden feature space of the deep learning model, while conversely with early integration we explored feature integration at the raw input feature level. In early integration, the original feed-forward deep learning model was applied where all three feature sets were merged together into one input vector. This architecture is the most simplistic yet it allows the model to extract hidden features using all input features already in the first layer right after the input layer. In multi-branch integration, the feature sets were fed into the model through separated branches and then the final individual dense layers were concatenated before calculating the final output. In consecutive integration, the feature sets are processed one after another and when the processing of a feature set is done, the last dense layer of that feature branch is concatenated with the next feature set into a new input layer (**Fig 1d**).

For all models, hyperparameter tuning was done by Bayesian optimization in the same fashion as in [5] and the optimization of the number of nodes and layers was performed after each input entry point (**Materials and methods**). All data integration experiments were carried out with the best performing data augmentation setting which varied across mutation downsampling rates (70% - 20x, 25% - 20x, 10% - 30x, 5% - 50x, 2% - 50x).

We found that including the 96 trinucleotide motifs and driver genes can improve the classification accuracy on the sparse somatic data. Early integration experiments resulted in the highest average accuracies overall, with an average accuracy of 0.65 compared to the data augmented baseline with 0.57 at 2%, with an average accuracy of 0.75 compared to the data augmented baseline with 0.69 at 5%, with an average accuracy of 0.82 compared to the data augmented baseline with 0.74 at 10% and with an average accuracy of 0.86 compared to the data augmented baseline with 0.84 at 25%, with the 70% sparse data experiment being the only exception where the average accuracy remained nearly the same at 0.884 compared to the data augmented baseline with 0.882 (**Fig 3a**). When assessing the performance of the best data integration model at cancer type level, for most cancer types a gradual accuracy improvement is apparent (**Fig 3b**). At 2% and 5% downsampling rates, the accuracy of the majority of cancer types shows the most notable improvement, which is especially appreciable in Kidney-RCC, Stomach-AdenoCA, Panc-Endocrine, Bone-Osteosarc and CNS-Medullo (**Fig 3b**). At other downsampling rates, where more SNVs are retained by default, certain cancer type accuracies slightly decrease (e.g.: Stomach-AdenoCA at 10% and Ovary-AdenoCA at 25%). In these cases the data augmentation might be insufficient to combat the increased dimensionality which moreover introduces additional noise caused by the additional features (**Fig 3b**). We conclude that utilizing additional, compact features derived from the raw SNV data can further aid the cancer type classification model, especially when only 2% or 5% of the somatic SNVs are retained in the sparse somatic samples.

### Assessment of top 1, 2 and 3 ranked predictions

**Fig 3.**
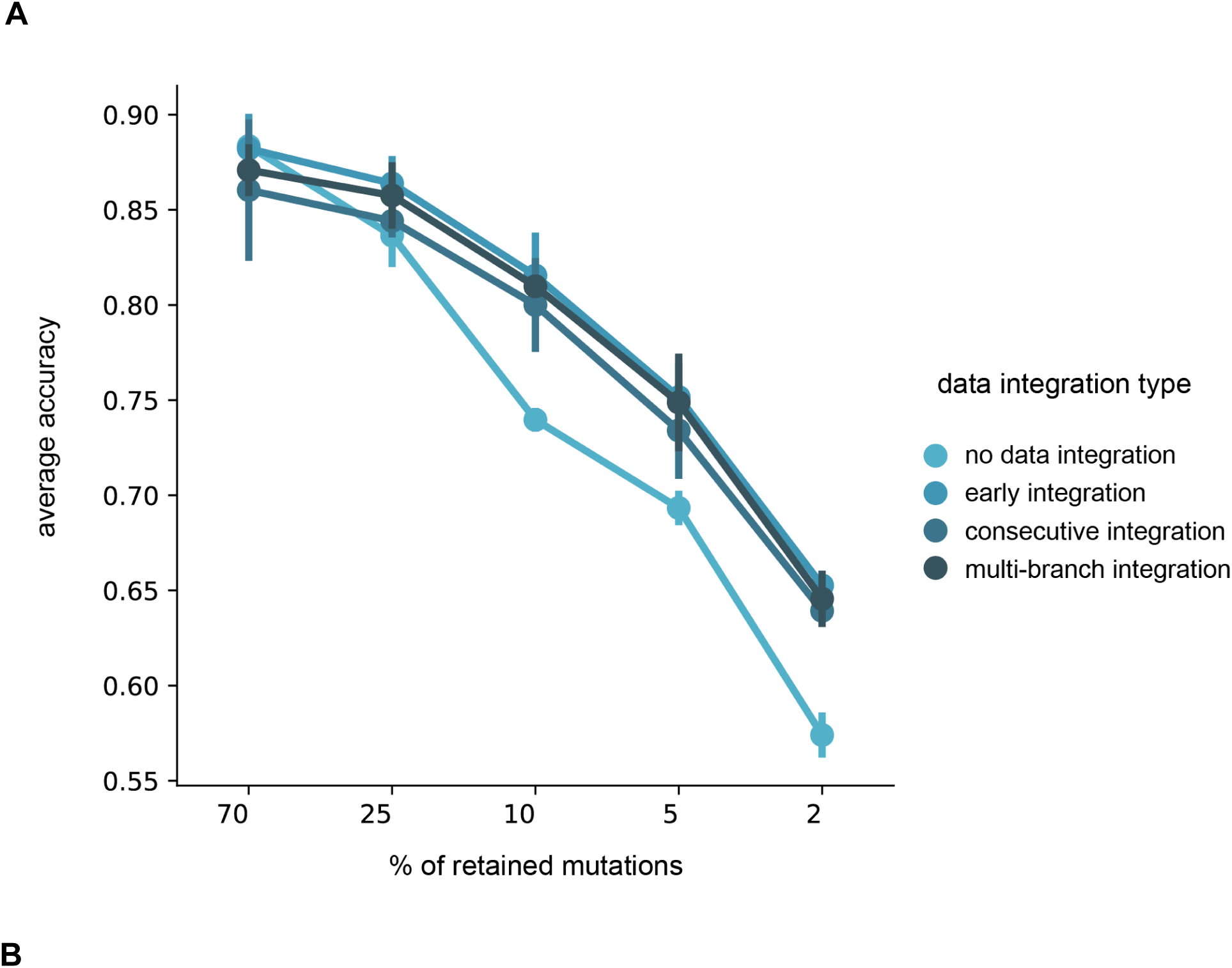
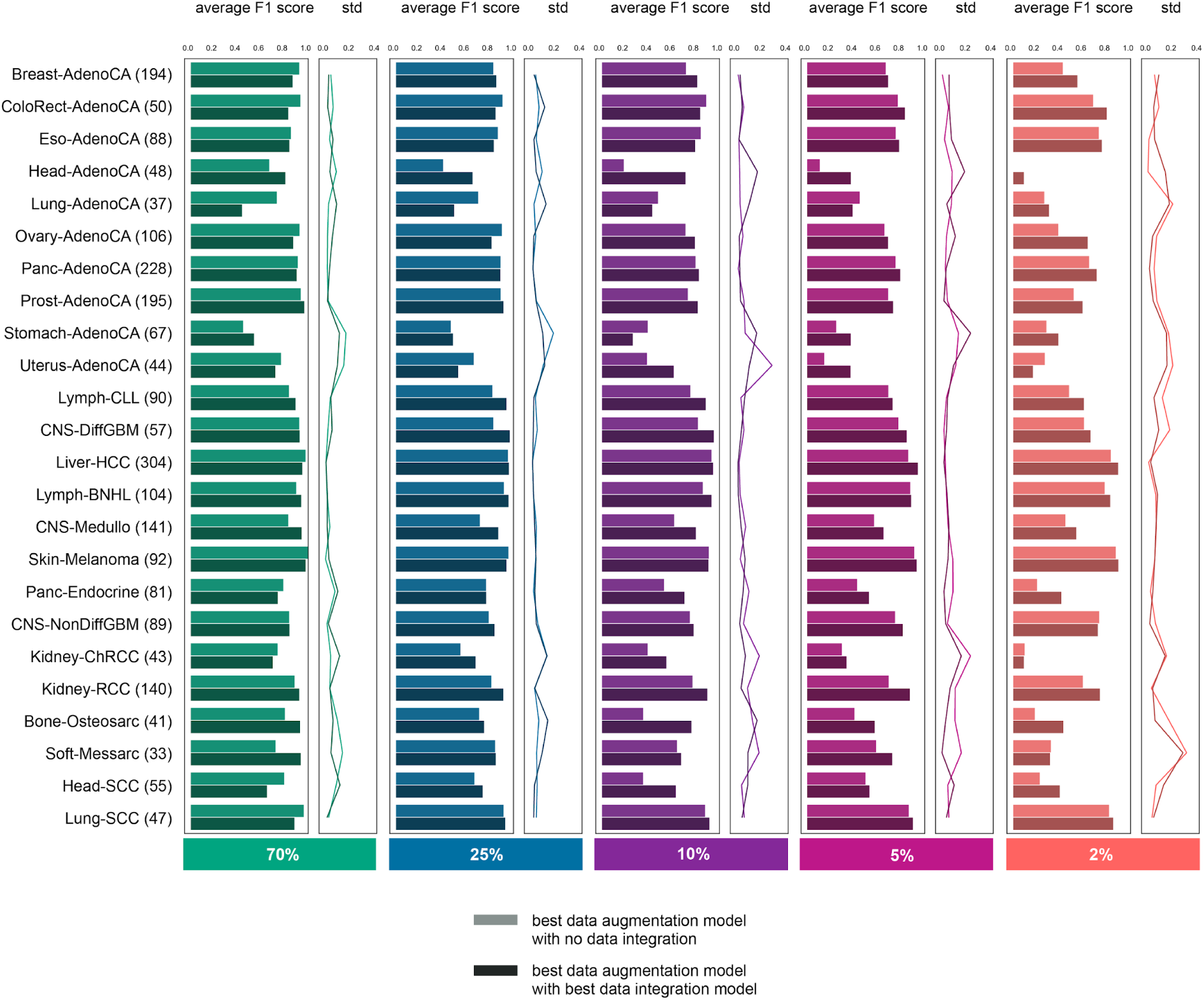
Classifier performance with data integration. **(A)** Overall effect of data integration at different mutation downsampling rates (70%, 25%, 10%, 5%, 2%). Average accuracy (average % of accurately classified samples in 3 folds) is shown at different downsampling rates with and without different applied data integration architectures **(B)** Classifier performance at different mutation downsampling rates with the best data integration architecture, expressed in F1-scores per cancer type (average and std is shown based on 3 folds). The best data integration architecture was selected based on average accuracy (Fig 3a) values, which was consistently the early data integration architecture across all downsampling rates. For comparison we used the best data augmentation model as the “model without data integration”.

In a diagnostic setting, where the aim of the cancer type classifier is to support the physician in making treatment decisions, it may be sufficient to provide a ranked list of predictions, rather than only the top prediction. This is especially true if a few predictions are close in terms of ranking based on the predicted cancer type probabilities (i.e. have very similar classification scores). These tumor types may have similar features, which could direct treatment decisions as well. We therefore also assessed the classifier accuracy when considering the top 1, 2 and 3 predictions of the model. When comparing the average accuracy of top 1 predictions (average ratio of correctly classified samples in stratified shuffle split cross-validation with three partitions) between different mutation downsampling rates with and without our model improvements, the final improved model achieves an average accuracy of 0.65 compared to an average accuracy of 0.41 with the original model at 2%, 0.75 compared to 0.56 at 5%, 0.82 compared to 0.63 at 10%, 0.86 compared to 0.75 at 25% and 0.88 compared to 0.83 at 70%, respectively (**Fig 4a**). When the original model trained on full somatic data is compared to different mutation downsampling rates with our model improvements in terms of average accuracy of the top 1 predictions, we found that the classifier performs slightly better at 70% downsampling rate than at 100% (**Fig 4a**). The underlying reason is likely that the noise introduced by the missing SNV calls functions as a regularizer at this rate and it improves on the generalization performance of the deep learning model during training and validation (at higher downsampling rates, this benefit clearly diminished since at those rates the lack of data is so severe that it decreases the overall performance). When the correct cancer type is identified from the top 3 predictions, the average accuracy is above 0.85 with our improvements, even at 2% downsampling rate (**Fig 4a**). When considering the top 2 and 3 predictions, the detection of all cancer types is improved, most prominently in case of Stomach-AdenoCA (**Fig 4b**).

### Feature importance assessment

**Fig 4.**
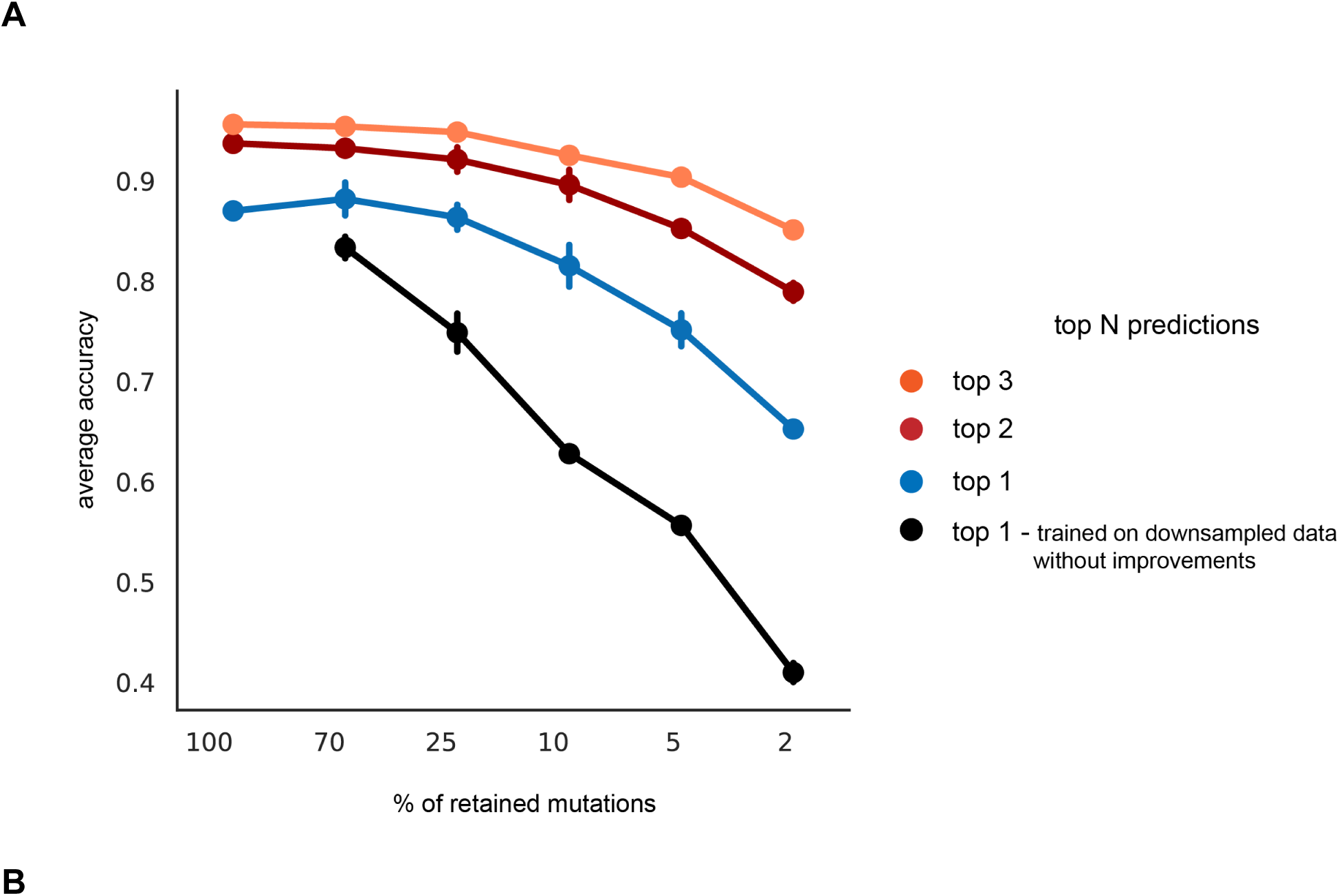
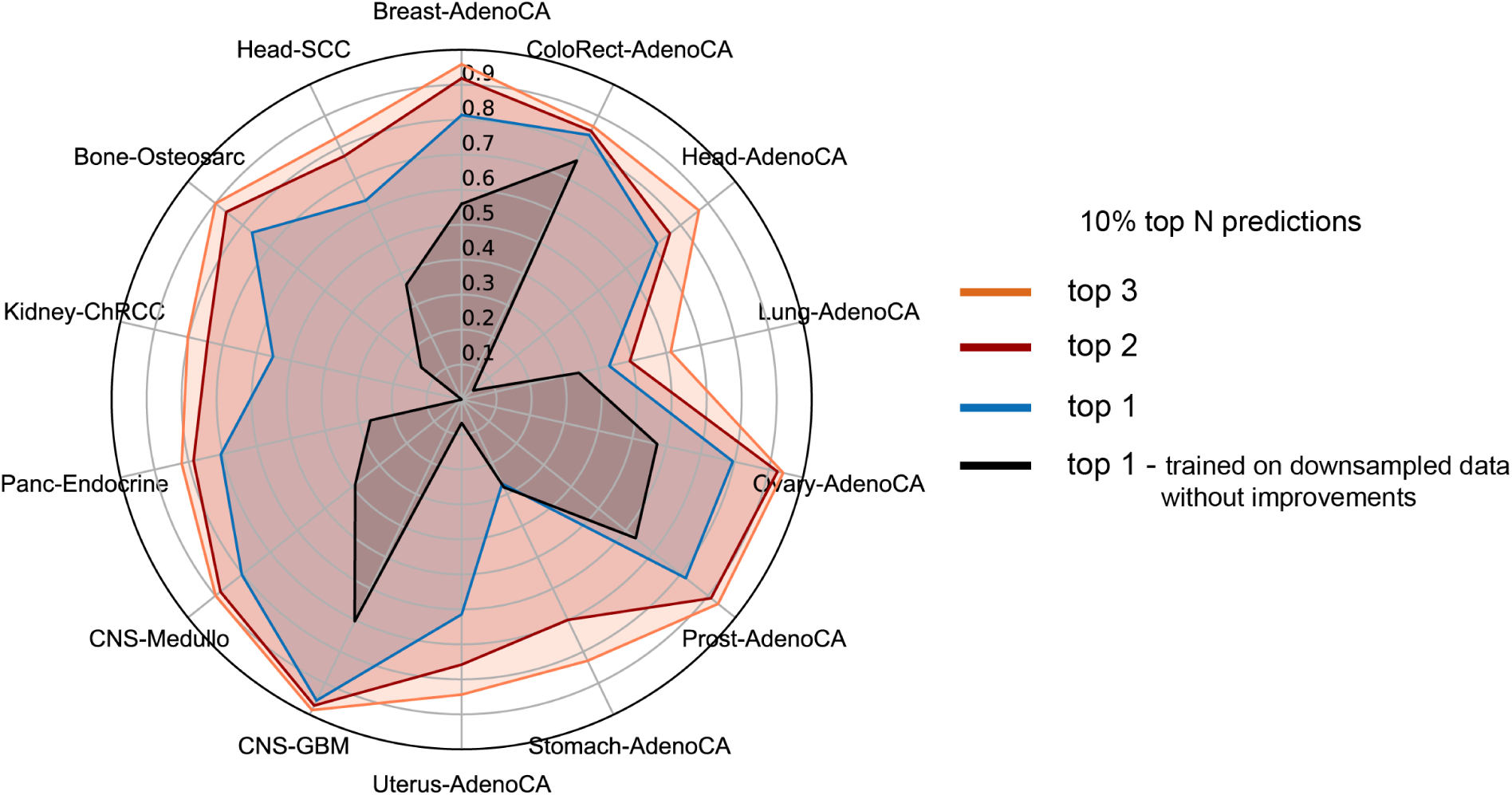
Classifier performance in predicting the top N labels. **(A)** Average accuracy (average % of accurately classified samples in 3 folds) when the correct cancer type is among top 1, 2 and 3 predictions. Average accuracy of the best models is shown at each mutation downsampling rate when identifying the correct tumour type among its top N-ranked predictions (N=1,2,3), error bars indicate standard deviation. **(B)** Classifier performance on somatic SNV data with 10% of retained mutations when the correct cancer type is among top 1, 2 and 3 predictions, compared to the baseline model. The axes represent the accuracy for each cancer type, expressed in average F1 scores (stratified shuffle split cross-validation with three partitions).

Interpretability is an important aspect in the application of machine learning, especially in the clinical field. By understanding why the model has made a certain prediction, the adoption of the approach in the clinic is facilitated as physicians can, to a certain extent, manually verify its decisions based on their own knowledge and experience. Model interpretation is usually done by feature importance assessment (or in other words feature attribution). However, defining feature importance is inherently problematic with deep learning models, due to the highly non-linear and complex nature of the models. As a result, along with the emergence of these models, solutions to assess feature attributions have also been developed. We aimed to use a straightforward method for our interpretation task and applied the so-called ‘integrated gradients’ method to obtain feature attributions [28] (see **Materials and methods**). In summary, the method takes the gradient of the output (probability of cancer type) with respect to the input feature values. In case of positive feature attribution, when the value of the feature was moving away from the baseline value then the probability of the given cancer type would increase, while in case of negative feature attribution, if the feature value moved closer to the baseline value then the target probability would increase [28]. Feature attribution is calculated per sample. To attain a feature importance within the context of a specific cancer type, feature attributions are averaged across the samples within the class.

First, we aimed to assess feature importances on the original, full SNV data and executed early data integration on it, in order to have feature importances calculated based on the full somatic profile of the PCAWG samples. The top 1 bin, top 2 trinucleotide and top 2 driver gene features were selected in each cancer type separately based on the ranked feature importances, and then the union of all top features across all cancer types (41 bins, 13 driver genes and 19 trinucleotides) were visualized (**Fig 5a**). Interestingly, in Skin-Melanoma, five of the positively attributed top trinucleotide features bear C>T nucleotide transition (CCC>CTC, CCT>CTT, GCA>GTA, TCC>TTC, TCT>TTT), while other positively attributed substitution types only appear in one or two trinucleotides. This is in line with the fact that this cancer type is dominated by C>T nucleotide transitions due to UV-light exposure [29]. *BCL2* has a high positive attribution in Lymph-BNHL and it was found to be frequently mutated in multiple lymphomas [30]. *ERBB4* is among the top attributed driver genes in Liver-HCC, which also corresponds to previous findings, where *ERBB4* was described as a suppressor in the development of Liver-HCC, suggesting that somatic mutations in *ERBB4* can function as drivers [31]. *ERBB4* also positively contributed to the prediction of Stomach-AdenoCA, Panc-AdenoCA, Eso-AdenoCA, Kidney-RCC and Lymph-CLL. *ERBB4* mutations were found in gastric carcinomas, albeit to a lesser extent than in colorectal and lung carcinomas [32], however to our knowledge, notable SNVs have not been reported in Lymph-CLL. Moreover, *ERBB4* was proposed as a tumor suppressor in Kidney-RCC [33], which might correlate with its positive feature attribution driven by increased somatic mutation density in *ERBB4* in Kidney-RCC. Importantly, we do not imply to draw direct connections with causality but rather aim to show that certain feature importances can capture meaningful underlying biological variation.

**Fig 5.**
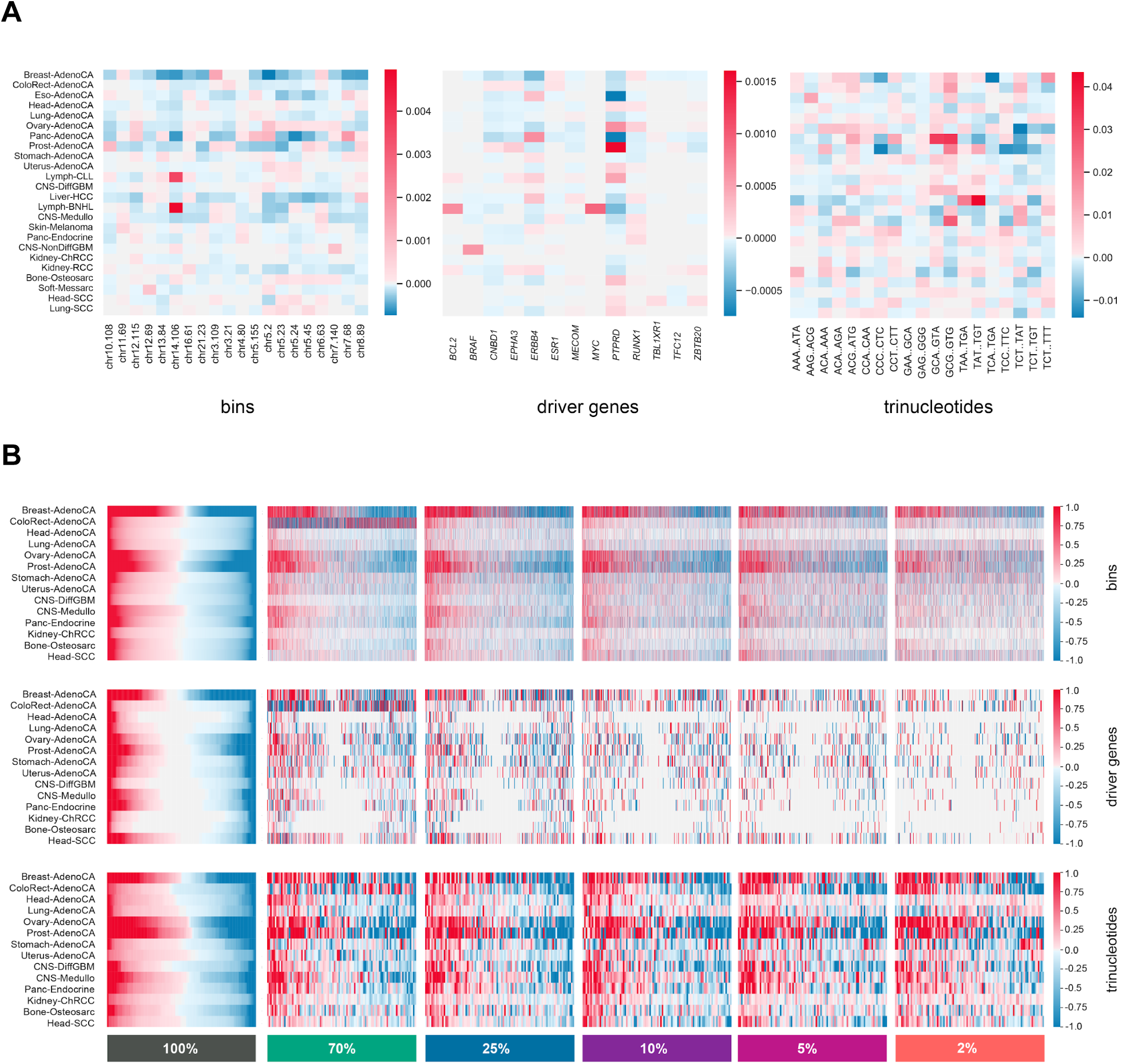
Feature importance assessment across all sparsity levels. **(A)** Top features in each feature type based on feature importance in full solid biopsy data. Top features were selected based on average feature importance ranking per cancer type in the test folds, resulting in 41 bins, 13 driver genes and 19 trinucleotides. The color scale represents negative (blue) and positive (red) feature importance values with zero (white) centering. **(B)** Feature importance patterns per feature type in full solid biopsy SNV data and sparse SNV data. Heatmaps of overall feature importance patterns were log-modulus transformed and scaled. Feature attributions were sorted per cancer type in the full solid biopsy SNV data and then the sparse data matrices were sorted to follow the same feature order per cancer type.

Overall changes in feature importance were compared between the original, full somatic data and sparse data settings in early data integration, as this architecture type resulted in the highest accuracy improvements in general. Our results show that the feature importance changes considerably in sparse data compared to the full SNV data (**Fig 5b**). The effect of SNV downsampling can be clearly seen when comparing feature importance profiles of the bin features in the baseline experiment with full somatic data to the experiments with sparse data, where many features have changed signs or have lessened or completely diminished importance due to sparsity. On the other hand, the 96 trinucleotide feature importances did not change drastically in sparse data, indicating that these features can still be harnessed more effectively by the classifier. This again validates our intuition, as we expected to see performance improvement on much sparser data with the inclusion of more SNV derived features.

## Discussion

Diagnosis of CUPs poses an important challenge in clinical practice. Utilizing the genome-wide somatic mutation profiles of ctDNA could offer a valuable approach to detect the cell of origin of cancer, but liquid biopsies bring about their own challenges as the inherently low concentration of ctDNA gives rise to sparse somatic mutation profiles. Our results show that both data augmentation and data integration can massively improve classifier performance, but the effect depends on the sparsity of mutational profiles. It appears that at less severe 70% and 25% downsampling, where more information is preserved from the original WGS samples by definition, the classifier cannot exploit the generated training samples of higher data augmentation levels (30-50x) and, hence, 20x data augmentation is sufficient to improve classifier performance. Data integration resulted in performance improvement in all sparse datasets except 70%, where the average accuracy remained nearly the same in comparison to the best data augmented model (0.884 and 0.882). When looking at the top 1 predictions with our improved model compared to the predictions made with the original model in all sparse datasets, the improved model achieves an average accuracy of 0.65 compared to an average accuracy of 0.41 with the original model at 2%, 0.75 compared to 0.56 at 5%, 0.82 compared to 0.63 at 10%, 0.86 compared to 0.75 at 25% and 0.88 compared to 0.83 at 70%, respectively, while in case the correct cancer type is identified from the top 3 predictions, our model reaches an average accuracy above 0.85, even at 2% and 5% downsampling rate (**Fig 4a**). This suggests that in case targeted treatments can be sufficiently selected based on the top 3 predicted cancer types, our classifier can aid treatment decisions even in extremely sparse data conditions.

In previous studies, the reported percentage of somatic mutations that can be detected in liquid biopsies compared to solid tissue biopsies in most advanced and metastatic cancers were 72% and 73.8% [13, 15], therefore our 70% sparse dataset was designated as an approximation of the reported values that can potentially reflect the sparsity in metastatic/CUP liquid biopsy data. As mentioned above, the final average accuracy at 70% is 0.88 compared to the baseline 0.83, and might indicate that this genome-wide SNV mutation information based approach can perform well in advanced and metastatic cancers. Interestingly, at 70%, the average accuracy has improved compared to the baseline model with full somatic SNV data (**Fig 4a**), which suggests that mutation downsampling can function as a regularizer at this rate, giving rise to improved performance. However, while retaining 70% of somatic SNVs provides a reasonable dataset to get an accuracy estimate for most tumor types in CUP/metastatic condition, this approximation might only be reliably applied for samples which are deep sequenced and cannot hold for samples with lower average coverage. Besides, certain cancer types such as glioma, medulloblastoma, bladder, renal and gastroesophageal cancer have been reported to have significantly lower ctDNA concentrations even in advanced disease [10]. For these tumor types, fewer variants will be detected in the blood and, therefore, lower downsampling rates provide a better estimation of classifier performance. With the improvements applied on the sparse dataset, a classification accuracy of 0.87 was achieved at 25% and 0.8 at 10% in CNS-Medullo (medulloblastoma), 0.97 accuracy was achieved at 25% and 0.95 at 10% in CNS-DiffGBM (diffuse glioma), 0.84 accuracy was achieved at 25% and 0.78 at 10% in CNS-DiffGBM (non-diffuse glioma), 0.68 accuracy was achieved at 25% and 0.55 at 10% in Kidney-ChRCC (renal cell carcinoma, distal tubule), 0.92 accuracy was achieved at 25% and 0.9 at 10% in Kidney-RCC (renal cell carcinoma, proximal tubule) and a classification accuracy of 0.84 was achieved at 25% and 0.79 at 10% in Eso-AdenoCA (esophageal adenocarcinoma).

The results presented here indicate that most of the problematic cancer types can still be classified with a reasonable accuracy even in very sparse somatic SNV data, which supports the diagnostic relevance of somatic SNV data derived from liquid biopsies. Moreover, since variants at low allele frequencies can be missed in solid biopsy samples at lower tumor purity levels, we hypothesise the applicability of our improvements to boost predictive performance on tissue biopsy samples, by means of predicting the cell of origin in tissue biopsy data with a classification model that was trained on sparse augmented data with data integration. This argument is bolstered by our finding where we compared the performance of the improved classifier on 70% downsampled data to the performance of the original model on the 100% SNV data, and the average accuracy of the 70% model was found to be improved compared to the baseline model (**Fig 4a**).

An important caveat in using somatic mutation profiles obtained from liquid biopsies is the presence of non-cancerous somatic mutations. According to a recent study, most of the non-cancerous cfDNA originates from white blood cells (WBC), therefore liquid biopsies contain a significant amount of somatic mutations as a result of clonal haematopoiesis in WBC [6, 13], which are indistinguishable from somatic mutations that arise in the cancer. This finding emphasises the importance of matched cfDNA WBC sequencing for accurate somatic variant interpretation in liquid biopsies. In our work, we only accounted for the sparse profile of somatic mutations that arise from ctDNA and considered the non-cancerous somatic SNVs and germline mutations to be already filtered out based on e.g. matched cfDNA WBC sequencing.

Despite the challenge of using sparse somatic data to identify the cell of origin of cancer, our results demonstrate that a deep learning classification model that uses multiple feature types and training set augmentation can help to harness the available sparse data in an efficient way. We conclude that somatic mutation density profiles obtained from liquid biopsies are suitable for the detection of the cell of origin of cancer, in particular for advanced and metastatic patients, and therefore they provide a valuable alternative to invasive solid tissue biopsies for CUP identification.

## Materials and methods

### Sparse data generation

Somatic SNV consensus calls of 2,374 primary tumors based on WGS with an average coverage of ∼30X were downloaded in VCF format from the ICGC Data Portal (http://dcc.icgc.org/releases/PCAWG/). The 2,374 primary tumor sample set was constructed from cancer types where at least 30 samples were available (**S1 Table**). The list of used PCAWG sample and donor identifiers can be found in **S2 Table**.

In order to model the sparse somatic mutation profile of ctDNA in liquid biopsy, we randomly subsampled 2%, 5%, 10%, 25% and 70% of the somatic SNVs from each primary cancer VCF file. The resulting random samples were then used to generate genome-wide SNV bin density, genome-wide trinucleotide and driver gene SNV density data for each sparse sample. A schematic overview of sparse sample generation can be seen in **Fig 1a**.

### Baseline/original model and model improvements

The original classifier [5] was downloaded from https://github.com/ICGC-TCGA-PanCancer/TumorType-WGS/tree/master/DNN-Model and retrained on the available PCAWG samples. In all consecutive steps, this model was used as a basis and further adjustments were applied on it in the data integration experiments. All additional models were implemented and trained in Tensorflow 1.10.0 and Keras 2.1.5. All code was written in Python 3.6.

We used shuffle split cross-validation with three splits, by randomly selecting 60% of the samples for training and 20-20% for validation and testing in each split round, respectively. Stratification was applied in each split, in order to distribute the cancer types according to the relative percentages in the training (60%), validation (20%) and test sets (20%). Hyperparameters were set by optimizing the performance on the validation set in each shuffle-split partition. The optimized hyperparameter set was the following: learning rate for Adam [34], L2-regularisation penalty (otherwise known as weight decay), dropout rate, the number of hidden layers, the number of neurons per hidden layer and activation function [5]. We used the same Bayesian method and library for hyperparameter optimization as in [5], where the ‘gp_minimize’ function from the scikit-optimise 0.7.4 library was applied [35]. Briefly, the models in each fold were trained using the Adam optimizer with a batch size of 32 for 50 epochs. Bias values were initialised as 0 and all other network weights were initialised using a glorot uniform distribution [36]. Each model was evaluated with 200 hyperparameter combinations that were obtained from the Bayesian optimization process (i.e., ‘gp_minimize’ function was called consecutively 200 times).

In the multi-branch and consecutive data integration experiments, the following hyperparameters were optimized separately for each feature type: weight decay, number of dense layers, number of dense nodes per layer, dropout and activation function. This meant that for the multi-branch experiments, hyperparameters were optimized per branch, while in the consecutive data integration experiments the hyperparameters were set separately after each input entry point. The final model architectures with data integration are shown in **S5-S19 Fig**. The final hyperparameter settings of each model are described in **S3 Table**.

### Accuracy calculation

Cancer type classification accuracies were calculated as F1-scores to take class imbalance into account, and averaged across the three shuffle split runs. This cross-validation setup was applied in all further experiments when comparing classification accuracy at cancer type level. Additionally, average model accuracy was calculated on the basis of correctly classified samples in all cancer types. F1-scores and confusion matrices were calculated with scikit-learn 0.23.2 [37].

### Data augmentation

N subsamples were generated from each sample, where N ∈ (10, 20, 30, 40, 50), N corresponds to the data augmentation level of the dataset which we refer to as ‘10x’, ‘20x’, ‘30x’, ‘40x’ and ‘50x’ (**S4 Table**). Sampled mutations were randomly selected for all N subsamples as described in ‘**Sparse data generation**’. If a subsampled sample was present in the training set, none of its other subsamples was part of the test/validation sets during the classification process.

### Data integration

The 96 trinucleotide motif features were generated on a per motif basis where all occurrences of a motif (e.g.: ACA>AGA) were aggregated, resulting in a summed value for the whole genome in each patient sample.

For the construction of driver gene features, we used the driver gene list released in one of the flagship papers of the latest the Pan-Cancer Atlas release [38]. SNVs were counted on gene regions from the start position of the first exon until the end position of the last exon.

### Feature importance assessment

The original published code (https://github.com/ankurtaly/Integrated-Gradients) from [28] was used to calculate the integrated gradients for the early integration models. Feature attributions were calculated in respect to the top predicted label in each sample. When calculating integrated gradients, first the gradient of the prediction output is calculated with respect to the feature values of the input in a range of interpolations between the original input (a vector containing all input features) and a baseline. In our case, zero baseline was used for all features. We calculated the feature attributions for each test split.

As this approach can be applied to explain feature attributions on a single sample level, the feature attributions were averaged from all test samples in each cancer type in order to get cancer type specific feature attributions. In order to aid data visualization when comparing the full sparse feature matrices, log-modulus transformation 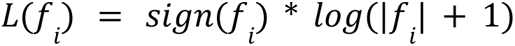 was applied on the raw feature importance values, where *f_i_* is the feature attribution value of feature *f* in cancer type *i*. Then we averaged the log-modulus transformed attributions of all test samples in each cancer type in order to get the transformed attributions per cancer type. Finally, scaling was applied to each feature matrix individually: 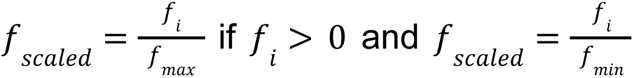 if 0 > *f_i_*. For visualization purposes, feature attributions were sorted per cancer type in the full solid biopsy SNV data and then the sparse data matrices were sorted based on the obtained feature order in the full solid biopsy SNV data, in order to follow the same feature order per cancer type in all matrices of a given feature type. Features that consistently had 0 attribution across all cancer types were excluded from visualization and are not displayed in the heatmaps.

### Code availability

Scripts that were used for classification, feature importance calculation and sparse augmented data generation are available at https://github.com/UMCUGenetics/cancer_type_classification_from_sparse_SNV_data, trained models are also available upon request. The code is distributed under the Apache Version 2.0 Open Source license (https://www.apache.org/ licenses/LICENSE-2.0).

### Data availability

The used PCAWG (https://doi.org/10.1038/s41586-020-1969-6) sample identifiers are listed in **S2 Table**. Aligned sequencing data, somatic and germline variant calls are available at https://dcc.icgc.org/releases/PCAWG. Further information regarding data access can be found at https://docs.icgc.org/pcawg/data/.

## Supporting information

**S1 Table. Sample distribution across cancer types in the used PCAWG dataset.**

**S2 Table. List of used PCAWG samples in the training, validation and test folds with donor identifiers and DCC project codes.**

**S3 Table. List of final optimized hyperparameters of all models.**

**S4 Table. Number of training samples with and without data augmentation in the 3 cross-validation folds.**

**S1 Fig. Classifier performance at different mutation downsampling rates (expressed in average F1 score and standard deviation of accuracy in folds) with the original model trained on the full (100%) data.** Test samples were generated by retaining 70%, 25%, 10%, 5% and 2% of somatic mutations from each original sample (mutation downsampling) while the training and validation set consisted of only original, full somatic SNV samples.

**S2 Fig. Classifier performance (expressed in average F1 score and standard deviation of accuracy in folds) at different mutation downsampling rates with models trained on sparse data sets.** Samples were generated by retaining 70%, 25%, 10%, 5% and 2% of somatic mutations from each original sample (mutation downsampling) for the training, validation and test sets.

**S3 Fig. Classifier performance (expressed in average F1 score and standard deviation of accuracy in folds) at different mutation downsampling rates with all the data augmentation setups.** Samples were generated by retaining 70%, 25%, 10%, 5% and 2% of somatic mutations from each original sample (mutation downsampling). Different data augmentation levels were assessed, see *Data augmentation* in **Materials and methods.**

**S4 Fig. Classifier performance (expressed in average F1 score and standard deviation of accuracy in folds) at different mutation downsampling rates with all the data integration setups on the best data augmentation model.** Samples were generated by retaining 70%, 25%, 10%, 5% and 2% of somatic mutations from each original sample (mutation downsampling). After performing data augmentation experiments, the best data augmentation setup was selected based on average accuracy (Supplementary Fig. 2) values: 70% - 20x, 25% - 20x, 10% - 30x, 5% - 50x, 2% - 50x.

**S5 Fig. Best model architecture selected based on hyperparameter search in early data integration on 2% downsampled sparse data. (A)** model in cross-validation fold 1 **(B)** model in cross-validation fold 2 **(C)** model in cross-validation fold 3

**S6 Fig. Best model architecture selected based on hyperparameter search in multi-branch data integration on 2% downsampled sparse data. (A) model in** cross-validation fold 1 **(B)** model in cross-validation fold 2 **(C)** model in cross-validation fold 3

**S7 Fig. Best model architecture selected based on hyperparameter search in consecutive data integration on 2% downsampled sparse data. (A)** model in cross-validation fold 1 **(B)** model in cross-validation fold 2 **(C)** model in cross-validation fold 3

**S8 Fig. Best model architecture selected based on hyperparameter search in early data integration on 5% downsampled sparse data. (A)** model in cross-validation fold 1 **(B)** model in cross-validation fold 2 **(C)** model in cross-validation fold 3

**S9 Fig. Best model architecture selected based on hyperparameter search in multi-branch data integration on 5% downsampled sparse data. (A)** model in cross-validation fold 1 **(B)** model in cross-validation fold 2 **(C)** model in cross-validation fold 3

**S10 Fig. Best model architecture selected based on hyperparameter search in consecutive data integration on 5% downsampled sparse data. (A)** model in cross-validation fold 1 **(B)** model in cross-validation fold 2 **(C)** model in cross-validation fold 3

**S11 Fig. Best model architecture selected based on hyperparameter search in early data integration on 10% downsampled sparse data. (A)** model in cross-validation fold 1 **(B)** model in cross-validation fold 2 **(C)** model in cross-validation fold 3

**S12 Fig. Best model architecture selected based on hyperparameter search in multi-branch data integration on 10% downsampled sparse data. (A)** model in cross-validation fold 1 **(B)** model in cross-validation fold 2 **(C)** model in cross-validation fold 3

**S13 Fig. Best model architecture selected based on hyperparameter search in consecutive data integration on 10% downsampled sparse data. (A)** model in cross-validation fold 1 **(B)** model in cross-validation fold 2 **(C)** model in cross-validation fold 3

**S14 Fig. Best model architecture selected based on hyperparameter search in early data integration on 25% downsampled sparse data. (A)** model in cross-validation fold 1 **(B)** model in cross-validation fold 2 **(C)** model in cross-validation fold 3

**S15 Fig. Best model architecture selected based on hyperparameter search in multi-branch data integration on 25% downsampled sparse data. (A)** model in cross-validation fold 1 **(B)** model in cross-validation fold 2 **(C)** model in cross-validation fold 3

**S16 Fig. Best model architecture selected based on hyperparameter search in consecutive data integration on 25% downsampled sparse data. (A)** model in cross-validation fold 1 **(B)** model in cross-validation fold 2 **(C)** model in cross-validation fold 3

**S17 Fig. Best model architecture selected based on hyperparameter search in early data integration on 70% downsampled sparse data. (A)** model in cross-validation fold 1 **(B)** model in cross-validation fold 2 **(C)** model in cross-validation fold 3

**S18 Fig. Best model architecture selected based on hyperparameter search in multi-branch data integration on 70% downsampled sparse data. (A)** model in cross-validation fold 1 **(B)** model in cross-validation fold 2 **(C)** model in cross-validation fold 3

**S19 Fig. Best model architecture selected based on hyperparameter search in consecutive data integration on 70% downsampled sparse data. (A)** model in cross-validation fold 1 **(B)** model in cross-validation fold 2 **(C)** model in cross-validation fold 3

## Acknowledgments

The results published here are based upon data generated by the International Cancer Genome Consortium (ICGC): https://dcc.icgc.org/pcawg.

## References

1. Goldie S.J., Chincarini G., Darido C. Targeted Therapy Against the Cell of Origin in Cutaneous Squamous Cell Carcinoma. Int. J. Mol. Sci. 2019;20:2201. https://doi.org/10.3390/ijms20092201.

2. Greco FA. Molecular diagnosis of the tissue of origin in cancer of unknown primary site: useful in patient management. Curr Treat Options Oncol. 2013 Dec;14(4):634–42. https://doi.org/10.1007/s11864-013-0257-1.

3. Pavlidis N, Khaled H, Gaafar R. A mini review on cancer of unknown primary site: a clinical puzzle for the oncologists. J Adv Res. 2015;6:375–82. https://doi.org/10.1016/j.jare.2014.11.007.

4. Salvadores M, Mas-Ponte D, Supek F. Passenger mutations accurately classify human tumors. PLoS Comput Biol. 2019;15(4): e1006953. https://doi.org/10.1371/journal.pcbi.1006953.

5. Jiao, W., Atwal, G., Polak, P. et al. A deep learning system accurately classifies primary and metastatic cancers using passenger mutation patterns. Nat Commun. 2020;11, 728. https://doi.org/10.1038/s41467-019-13825-8.

6. Overman MJ, Modak J, Kopetz S, Murthy R, Yao JC, Hicks ME, Abbruzzese JL, Tam AL. Use of research biopsies in clinical trials: are risks and benefits adequately discussed? J Clin Oncol. 2013 Jan 1;31(1):17–22. https://doi.org/10.1200/JCO.2012.43.1718.

7. Cohen JD, Li L, Wang Y, Thoburn C, Afsari B, Danilova L, Douville C, Javed AA, Wong F, Mattox A, Hruban RH, Wolfgang CL, Goggins MG, Dal Molin M, Wang TL, Roden R, Klein AP, Ptak J, Dobbyn L, Schaefer J, Silliman N, Popoli M, Vogelstein JT, Browne JD, Schoen RE, Brand RE, Tie J, Gibbs P, Wong HL, Mansfield AS, Jen J, Hanash SM, Falconi M, Allen PJ, Zhou S, Bettegowda C, Diaz LA Jr, Tomasetti C, Kinzler KW, Vogelstein B, Lennon AM, Papadopoulos N. Detection and localization of surgically resectable cancers with a multi-analyte blood test. Science. 2018 Feb 23;359(6378):926–930. https://doi.org/10.1126/science.aar3247.

8. Sung J.S., Chong H.Y., Kwon N.-J., Kim H.M., Lee J.W., Kim B., Lee S.B., Park C.W., Choi J.Y., Chang W.J., et al. Detection of somatic variants and EGFR mutations in cell-free DNA from non-small cell lung cancer patients by ultra-deep sequencing using the ion ampliseq cancer hotspot panel and droplet digital polymerase chain reaction. Oncotarget. 2017;8:106901. https://doi.org/10.18632/oncotarget.22456.

9. Iwahashi, N., Sakai, K., Noguchi, T. et al. Liquid biopsy-based comprehensive gene mutation profiling for gynecological cancer using CAncer Personalized Profiling by deep Sequencing. Sci Rep. 2019;9, 10426. https://doi.org/10.1038/s41598-019-47030-w

10. Bettegowda C, Sausen M, Leary RJ, Kinde I, Wang Y, Agrawal N, Bartlett BR, Wang H, Luber B, Alani RM, Antonarakis ES, Azad NS, Bardelli A, Brem H, Cameron JL, Lee CC, Fecher LA, Gallia GL, Gibbs P, Le D, Giuntoli RL, Goggins M, Hogarty MD, Holdhoff M, Hong SM, Jiao Y, Juhl HH, Kim JJ, Siravegna G, Laheru DA, Lauricella C, Lim M, Lipson EJ, Marie SK, Netto GJ, Oliner KS, Olivi A, Olsson L, Riggins GJ, Sartore-Bianchi A, Schmidt K, Shih lM, Oba-Shinjo SM, Siena S, Theodorescu D, Tie J, Harkins TT, Veronese S, Wang TL, Weingart JD, Wolfgang CL, Wood LD, Xing D, Hruban RH, Wu J, Allen PJ, Schmidt CM, Choti MA, Velculescu VE, Kinzler KW, Vogelstein B, Papadopoulos N, Diaz LA Jr. Detection of circulating tumor DNA in early- and late-stage human malignancies. Sci Transl Med. 2014 Feb 19;6(224):224ra24. https://doi.org/10.1126/scitranslmed.3007094.

11. Haber DA, Velculescu VE. Blood-based analyses of cancer: circulating tumor cells and circulating tumor DNA. Cancer Discov. 2014 Jun;4(6):650–61. https://doi.org/10.1158/2159-8290.CD-13-1014.

12. Higgins MJ, Jelovac D, Barnathan E, Blair B, Slater S, Powers P, Zorzi J, Jeter SC, Oliver GR, Fetting J, Emens L, Riley C, Stearns V, Diehl F, Angenendt P, Huang P, Cope L, Argani P, Murphy KM, Bachman KE, Greshock J, Wolff AC, Park BH. Detection of tumor PIK3CA status in metastatic breast cancer using peripheral blood. Clin Cancer Res. 2012 Jun 15;18(12):3462–9. https://doi.org/10.1158/1078-0432.CCR-11-2696.

13. Razavi P, Li BT, Brown DN, Jung B, Hubbell E, Shen R, Abida W, Juluru K, De Bruijn I, Hou C, Venn O, Lim R, Anand A, Maddala T, Gnerre S, Vijaya Satya R, Liu Q, Shen L, Eattock N, Yue J, Blocker AW, Lee M, Sehnert A, Xu H, Hall MP, Santiago-Zayas A, Novotny WF, Isbell JM, Rusch VW, Plitas G, Heerdt AS, Ladanyi M, Hyman DM, Jones DR, Morrow M, Riely GJ, Scher HI, Rudin CM, Robson ME, Diaz LA Jr, Solit DB, Aravanis AM, Reis-Filho JS. High-intensity sequencing reveals the sources of plasma circulating cell-free DNA variants. Nat Med. 2019 Dec;25(12):1928–1937. https://doi.org/10.1038/s41591-019-0652-7.

14. Adalsteinsson, V.A., Ha, G., Freeman, S.S. et al. Scalable whole-exome sequencing of cell-free DNA reveals high concordance with metastatic tumors. Nat Commun 8, 1324 (2017). https://doi.org/10.1038/s41467-017-00965-y.

15. Jiang J, Adams HP, Yao L, Yaung S, Lal P, Balasubramanyam A, Fuhlbrück F, Tikoo N, Lovejoy AF, Froehler S, Fang LT, Achenbach HJ, Floegel R, Krügel R, Palma JF. Concordance of Genomic Alterations by Next-Generation Sequencing in Tumor Tissue versus Cell-Free DNA in Stage I-IV Non-Small Cell Lung Cancer. J Mol Diagn. 2020 Feb;22(2):228–235. https://doi.org/10.1016/j.jmoldx.2019.10.013.

16. McCabe, M.J., Gauthier, ME.A., Chan, CL. et al. Development and validation of a targeted gene sequencing panel for application to disparate cancers. Sci Rep. 2019;9, 17052. https://doi.org/10.1038/s41598-019-52000-3.

17. Zviran A, Schulman RC, Shah M, Hill STK, Deochand S, Khamnei CC, Maloney D, Patel K, Liao W, Widman AJ, Wong P, Callahan MK, Ha G, Reed S, Rotem D, Frederick D, Sharova T, Miao B, Kim T, Gydush G, Rhoades J, Huang KY, Omans ND, Bolan PO, Lipsky AH, Ang C, Malbari M, Spinelli CF, Kazancioglu S, Runnels AM, Fennessey S, Stolte C, Gaiti F, Inghirami GG, Adalsteinsson V, Houck-Loomis B, Ishii J, Wolchok JD, Boland G, Robine N, Altorki NK, Landau DA. Genome-wide cell-free DNA mutational integration enables ultra-sensitive cancer monitoring. Nat Med. 2020 Jul;26(7):1114–1124. https://doi.org/10.1038/s41591-020-0915-3.

18. The ICGC/TCGA Pan-Cancer Analysis of Whole Genomes Consortium., Campbell, P.J., Getz, G. et al. Pan-cancer analysis of whole genomes. Nature. 2020;578, 82–93. https://doi.org/10.1038/s41586-020-1969-6.

19. Alexandrov LB, Nik-Zainal S, Wedge DC, Aparicio SA, Behjati S, Biankin AV, Bignell GR, Bolli N, Borg A, Børresen-Dale AL, Boyault S, Burkhardt B, Butler AP, Caldas C, Davies HR, Desmedt C, Eils R, Eyfjörd JE, Foekens JA, Greaves M, Hosoda F, Hutter B, Ilicic T, Imbeaud S, Imielinski M, Jäger N, Jones DT, Jones D, Knappskog S, Kool M, Lakhani SR, López-Otín C, Martin S, Munshi NC, Nakamura H, Northcott PA, Pajic M, Papaemmanuil E, Paradiso A, Pearson JV, Puente XS, Raine K, Ramakrishna M, Richardson AL, Richter J, Rosenstiel P, Schlesner M, Schumacher TN, Span PN, Teague JW, Totoki Y, Tutt AN, Valdés-Mas R, van Buuren MM, van ’t Veer L, Vincent-Salomon A, Waddell N, Yates LR; Australian Pancreatic Cancer Genome Initiative; ICGC Breast Cancer Consortium; ICGC MMML-Seq Consortium; ICGC PedBrain, Zucman-Rossi J, Futreal PA, McDermott U, Lichter P, Meyerson M, Grimmond SM, Siebert R, Campo E, Shibata T, Pfister SM, Campbell PJ, Stratton MR. Signatures of mutational processes in human cancer. Nature. 2013 Aug 22;500(7463):415–21. https://doi.org/10.1038/nature12477.

20. Hanahan D, Weinberg RA. Hallmarks of cancer: the next generation. Cell. 2011 Mar 4;144(5):646–74. https://doi.org/10.1016/j.cell.2011.02.013.

21. Schaefer, M., Serrano, L. Cell type-specific properties and environment shape tissue specificity of cancer genes. Sci Rep. 2016;6, 20707. https://doi.org/10.1038/srep20707.

22. Shorten, C., Khoshgoftaar, T.M. A survey on Image Data Augmentation for Deep Learning. J Big Data. 2019:6, 60. https://doi.org/10.1186/s40537-019-0197-0.

23. Zhun Z, Liang Z, Guoliang K, Shaozi L, Yi Y. Random erasing data augmentation. arXiv:1708.04896v2 [Preprint] 2017 [cited 2021 Mar 3]. Database: arXiv [Internet]. Available from: https://arxiv.org/abs/1708.04896

24. Alex Krizhevsky, Ilya Sutskever, and Geoffrey E. Hinton. 2017. ImageNet classification with deep convolutional neural networks. Commun ACM. 2017 June;60, 6, 84–90. https://doi.org/10.1145/3065386.

25. Cecilia S, Michael JD. Improved mixed-example data augmentation. arXiv:1805.11272v4 [Preprint] 2018 [cited 2021 Mar 3]. Database: arXiv [Internet]. Available from: https://arxiv.org/abs/1805.11272

26. Hiroshi Inoue. Data Augmentation by Pairing Samples for Images Classification. arXiv:1801.02929v2 [Preprint] 2018 [cited 2021 Mar 3]. Database: arXiv [Internet]. Available from: https://arxiv.org/abs/1801.02929

27. Joel Hestness, Sharan Narang, Newsha Ardalani, Gregory Diamos, Heewoo Jun, Hassan Kianinejad, Md. Mostofa Ali Patwary, Yang Yang, Yanqi Zhou. Deep Learning Scaling is Predictable, Empirically. arXiv:1712.00409v1 [Preprint] 2017 [cited 2021 Mar 3]. Database: arXiv [Internet]. Available from:

28. Mukund Sundararajan, Ankur Taly and Qiqi Yan. Axiomatic Attribution for Deep Networks. arXiv:1703.01365v2 [Preprint] 2017 [cited 2021 Mar 3]. Database: arXiv [Internet]. Available from: https://arxiv.org/abs/1703.01365

29. Hodis E, Watson IR, Kryukov GV, Arold ST, Imielinski M, Theurillat JP, Nickerson E, Auclair D, Li L, Place C, Dicara D, Ramos AH, Lawrence MS, Cibulskis K, Sivachenko A, Voet D, Saksena G, Stransky N, Onofrio RC, Winckler W, Ardlie K, Wagle N, Wargo J, Chong K, Morton DL, Stemke-Hale K, Chen G, Noble M, Meyerson M, Ladbury JE, Davies MA, Gershenwald JE, Wagner SN, Hoon DS, Schadendorf D, Lander ES, Gabriel SB, Getz G, Garraway LA, Chin L. A landscape of driver mutations in melanoma. Cell. 2012 Jul 20;150(2):251–63. https://doi.org/10.1016/j.cell.2012.06.024.

30. Schuetz, J., Johnson, N., Morin, R. et al. BCL2 mutations in diffuse large B-cell lymphoma. Leukemia. 2012;26, 1383–1390. https://doi.org/10.1038/leu.2011.378.

31. Yao Liu, Liming Song, Hengli Ni, Lina Sun, Weijuan Jiao, Lin Chen, Qun Zhou, Tong Shen, Hongxia Cui, Tianming Gao, Jianming Li. ERBB4 acts as a suppressor in the development of hepatocellular carcinoma. Carcinogenesis. 2017 April;Volume 38, Issue 4,465–473. https://doi.org/10.1093/carcin/bgx017.

32. Soung YH, Lee JW, Kim SY, Wang YP, Jo KH, Moon SW, Park WS, Nam SW, Lee JY, Yoo NJ, Lee SH. Somatic mutations of the ERBB4 kinase domain in human cancers. Int J Cancer. 2006 Mar 15;118(6):1426–9. https://doi.org/10.1002/ijc.21507.

33. Thomasson M, Hedman H, Junttila TT, Elenius K, Ljungberg B, Henriksson R. ErbB4 is downregulated in renal cell carcinoma--a quantitative RT-PCR and immunohistochemical analysis of the epidermal growth factor receptor family. Acta Oncol. 2004;43(5):453–9. https://doi.org/10.1080/02841860410028574.

34. Kingma, D. P. & Ba, J. Adam: a method for stochastic optimization. arXiv:1412.6980v9 [Preprint] 2014 [cited 2021 Mar 3]. Database: arXiv [Internet]. Available from: https://arxiv.org/abs/1412.6980

35. Head, T., et al. scikit-optimize/scikit-optimize. 2020 Sept, https://doi.org/10.5281/zenodo.1157319.

36. Glorot, X.; Bengio, Y. Understanding the difficulty of training deep feedforward neural networks. In Proceedings of the Thirteenth International Conference on Artificial Intelligence and Statistics; Teh, Y.W., Titterington, M., Eds.; Chia Laguna Resort, PMLR: Sardinia, Italy, 2010; Volume 9, pp. 249–256.

37. Pedregosa et al., Scikit-learn: Machine Learning in Python. JMLR 2011;12, pp. 2825–2830. Available at: https://jmlr.csail.mit.edu/papers/v12/pedregosa11a.html

38. Bailey MH, Tokheim C, Porta-Pardo E, Sengupta S, Bertrand D, Weerasinghe A, Colaprico A, Wendl MC, Kim J, Reardon B, Ng PK, Jeong KJ, Cao S, Wang Z, Gao J, Gao Q, Wang F, Liu EM, Mularoni L, Rubio-Perez C, Nagarajan N, Cortés-Ciriano I, Zhou DC, Liang WW, Hess JM, Yellapantula VD, Tamborero D, Gonzalez-Perez A, Suphavilai C, Ko JY, Khurana E, Park PJ, Van Allen EM, Liang H; MC3 Working Group; Cancer Genome Atlas Research Network, Lawrence MS, Godzik A, Lopez-Bigas N, Stuart J, Wheeler D, Getz G, Chen K, Lazar AJ, Mills GB, Karchin R, Ding L. Comprehensive Characterization of Cancer Driver Genes and Mutations. Cell. 2018 Apr 5;173(2):371–385.e18. https://doi.org/10.1016/j.cell.2018.02.060.

